# Adaptive time course of the skeletal muscle proteome during programmed resistance training in rats

**DOI:** 10.1101/2025.02.17.633647

**Authors:** Connor A Stead, Stuart J Hesketh, Aaron C.Q Thomas, Mark R Viggars, Hazel Sutherland, Jonathan C Jarvis, Jatin G Burniston

## Abstract

Resistance training (RT) promotes muscle protein accretion and myofiber hypertrophy, driven by dynamic processes of protein synthesis and degradation. While molecular studies have focused on acute signalling or long-term hypertrophy and strength gains, a critical gap remains in understanding the intermediate processes of muscle adaptation. Acute signalling does not always correlate directly with long-term outcomes, highlighting the need for a time-course analysis of protein abundance and turnover rates. To address this, we utilised deuterium oxide labelling and peptide mass spectrometry to quantify absolute protein content and synthesis rates in skeletal muscle. A daily programmed resistance training regimen was applied to the rat tibialis anterior (TA) via electrical stimulation of the left hind limb for 10, 20, and 30 days (5 sets of 10 repetitions daily). Muscle samples from stimulated (Stim) and contralateral control (Ctrl) limbs were analysed, quantifying 658 protein abundances and 215 protein synthesis rates. Unsupervised temporal clustering of protein responses revealed distinct phases of muscle adaptation, with early (0-10 days) and mid (10-20 days) responses driven by differential protein accretion rates in ribosomal and mitochondrial networks, respectively. These findings suggest that subsets of proteins exhibit distinct adaptation timelines due to variations in translation and/or degradation rates. A deeper understanding of these temporal shifts could improve strategies for optimising muscle growth and functional adaptation to resistance training.

## Introduction

Resistance training (RT) is well recognised as a powerful stimulus for skeletal muscle adaptation, driving increases in the mass and strength of skeletal muscle that protect against age-related chronic diseases (McLeod et al., 2019). Gains in muscle mass are underpinned by protein accretion and may be accompanied by qualitative changes to the protein composition (i.e. proteome) of muscle that results in changes to muscle function, including speed of contraction and resistance to fatigue. Changes to the muscle proteome occur when the balance between the synthesis and degradation of individual proteins change (Hesketh et al., 2021). The rates of synthesis and degradation are directed by intracellular mechanisms that integrate the mechanical signals with other information (e.g. energy balance, endocrine state, developmental stage, immune function etc.) to appropriately allocate cellular resources to muscle growth (Roberts et al., 2023). There is strong interest in optimising the positive outcomes of muscle hypertrophy, fatigue resistance and metabolism, to enhance athletic performance, prevent chronic disease, and mitigate age-related functional decline. Nevertheless, knowledge on the mechanisms of adaptation is incomplete and this limits our ability to optimise the process of muscle adaptation to suit specific outcomes and populations.

Muscle hypertrophy is the most common objective of RT and substantial research effort has been dedicated to optimising muscle protein synthetic responses in humans (Davies et al., 2024). The fractional synthetic rate (FSR) of mixed myofibrillar proteins is known to double during the first 3-h period following a bout of resistance exercise (Phillips et al., 1997). However, the magnitude of the synthetic response measured by short-term infusion of amino acid tracers greatly exceeds or ‘over predicts’ the gains in muscle mass accrued over weeks of RT, in part, because muscle protein degradation is also elevated during the hours following resistance exercise (Phillips et al., 1997). When protein synthesis is measured over a timespan of weeks, using deuterium oxide (D_2_O), RT is associated with an ∼18.5 % increase in FSR (Brook et al., 2015) that reflects the net contributions of synthesis and degradation and correlates positively with gains in muscle size. D_2_O labelling can be readily combined with proteomic techniques to investigate the synthesis rate of individual muscle proteins (Burniston, 2019) and has been used to study muscle responses to RT in humans (Camera et al., 2017, Murphy et al., 2018) and selective androgen receptor modulators (SARM) in rats (Shankaran et al., 2016). In rat muscle, the FSR of 34 proteins including myofibrillar proteins and glycolytic enzymes, increased in a dose-dependent manner during the first week of SARM treatment, and 70 % (65 out of 94 proteins studied) of proteins exhibited positive correlations between the magnitude of increase in FSR during the first week and gains in muscle mass after 28 days of SARM treatment (Shankaran et al., 2016).

Correlative analyses between early protein FSR responses and subsequent gains in muscle mass cannot demonstrate the link between synthesis, degradation and changes in abundance on a protein-by-protein basis. Instead, dynamic proteomic profiling (DPP), encompassing measurements on both the abundance and synthesis rate of individual proteins, is required (Hesketh et al., 2021). When studied using DPP, the response of human muscle to RT (Camera et al., 2017) or high-intensity interval training (Srisawat et al., 2023) includes proteins that increase in FSR without exhibiting a change in abundance (i.e. turnover rate is increased), and proteins that decrease in abundance despite exhibiting an increase in FSR. So, there must also be changes in protein degradation. In short, the efficacy of RT protocols to accrue muscle mass cannot be assessed by the magnitude of the protein synthetic response alone.

In laboratory animals (Hesketh et al., 2020), and cell cultures (Brown et al., 2021), the <u>A</u>bsolute <u>D</u>ynamic <u>P</u>rofiling <u>T</u>echnique for <u>Proteo</u>mics (Proteo-ADPT) offers the opportunity to include measurements on total protein content and thereby report absolute rates (e.g. µg/d) of synthesis and degradation on a protein-by-protein basis. Using Proteo-ADPT, (Hesketh et al., 2020) reports the transformation of rat fast-twitch muscle by chronic low-frequency stimulation (CLFS) involves approximately equal contributions from changes in protein degradation and synthesis. In some instances, changes in degradation rate primarily drive gains in protein abundance, while some decreases in protein abundance occur primarily via a decrease in synthesis (Hesketh et al., 2020). CLFS is associated with 50 % reduction in muscle mass over 4 weeks (Hesketh et al., 2020) so, herein, we sought to investigate the role of changes in protein dynamics during gains in muscle mass using unilatetal programmed resistance training (PRT). PRT involves programmed co-contraction of rat lower limb muscles in vivo that leads to significant (∼15 %) gains in muscle mass during a 30-d experimental period (Schmoll et al., 2018, Viggars et al., 2023). In the current work, exercised (i.e. PRT) and contralateral control muscles were studied during early (day 0-10), mid (day 10-20) and late (day 20-30) periods of adaptation. Our findings highlight substantial ‘over synthesis’ of protein during the first 10 days of PRT and we discovered different temporal patterns of adaptation between ribosomal and mitochondrial proteins.

## Materials and Methods

### Animals and experimental design

Experimental procedures were conducted under the auspices of the British Home Office Animals (Scientific Procedures) Act 1986 (License number: PA693D221). Three month old, male Wistar rats (412 ± 69 g body weight; n = 16) were bred in-house in a conventional colony, housed in controlled conditions of 20 °C, 45 % relative humidity, under a 12 h light (0600–1800 hours) and 12 h dark cycle, water and food were available ad libitum. Animals were assigned to one of four groups (n = 4 in each), including a sham-operated control group and three experimental groups that received deuterium oxide (D_2_O; Sigma-Aldrich, St. Louis, MO) for either 10 d, 20 d or 30 d duration. D_2_O was initiated by an intraperitoneal bolus of 10 µL 99 % D_2_O-saline/ g body weight and maintained by supplementing the animals’ drinking water with 5 % (v/v) D_2_O, which was refreshed daily.

Unilateral programmed resistance training (PRT) was used to simulate high-intensity resistance training using an implanted device. Electrical nerve stimulation intended to induce hypertrophy by loaded contractions of the tibialis anterior (TA) was conducted as described previously by our group (Schmoll et al., 2018). Buprenorphine (Temgesic, Indivior, Slough, UK) at a dose of 0.05 mg/kg^-1^ body mass, was administered preoperatively for analgesia and surgery was performed with aseptic precautions. Anaesthesia was induced using a gaseous mixture of 4 % isoflurane in medical oxygen and was then adjusted to 1-2 % isoflurane to maintain an adequate plane of surgical anaesthesia. Electrodes were placed underneath the common peroneal nerve (cathode) and near to the tibial nerve (anode), to exploit the different activation thresholds for anodic and cathodic stimulation. The lower threshold of cathodic stimulation recruits all axons of the peroneal nerve so the dorsiflexors, including the TA, are maximally activated. An amplitude was set that provided sufficient additional activation of the plantar-flexors via the tibial nerve to resist the action of the dorsi-flexors so that the ankle joint angle did not decrease. Force produced by the stronger plantar-flexor muscles was transmitted via the ankle and was used to achieve auxotonic contractions of the TA. The 0-day time point represents the sham-operated control group that were implanted with inactive stimulators and then killed after the 1-week recovery period. The remaining animals had the stimulation patterns implemented remotely one week after the implant operation using the Mini-VStim-App installed on a standard Android driven tablet computer (Xperia Tablet Z, Sony Corporation, Tokyo, Japan). The tablet computer connected via Bluetooth connection to a Mini-VStim programming device which communicated with the pulse generator via a radio frequency link. Once programmed the stimulator operated autonomously to provide daily exercise sessions in one limb, and the animals were housed in their normal cages and monitored daily.

The stimulation pattern consisted of a daily ‘warm-up’ phase of 40 twitches at 4 Hz over a duration of 10 seconds, which immediately preceded the resistance training programme. The resistance training stimulus used 5 sets of 10 repetitions. Each repetition consisted of a 2 s fused near-maximal tetanic contraction at a stimulation frequency of 100 Hz. Two seconds recovery was allowed between repetitions and 2.5 minutes of rest was allowed between sets. One hour after daily stimulation on days 10, 20, and 30, groups of n=4 animals were killed humanely in a rising concentration of CO_2_ followed by cervical dislocation. Plasma samples were obtained by cardiac puncture immediately after death and TA muscles, from the left stimulated (Stim) and the right non-stimulated (Ctrl) limb were isolated. Each muscle was cleaned of fat and connective tissue then weighed before being frozen in liquid nitrogen and stored at -80 °C pending further analysis.

### Muscle processing

Muscles were fractionated into myofibrillar and soluble fractions according to Hesketh et al, (Hesketh et al., 2020). Samples were pulverized under liquid nitrogen using a mortar and pestle. An analytical balance was used to accurately record an aliquot (∼100 mg) of muscle powder, which was then mechanically homogenized on ice in 10 volumes of 1 % Triton X-100, 50 mM Tris pH 7.4 including phosphatase inhibitor and complete protease inhibitor cocktails (Roche, Indianapolis, USA). Homogenates were incubated on ice for 15 min, then centrifuged at 1000 x g, 4 °C for 5 min. Supernatants containing soluble proteins were decanted and stored on ice while the myofibrillar pellet was washed by resuspension in 0.5 ml of homogenization buffer and then centrifuged at 1000 x g, 4 °C for 5 min. The washed myofibrillar pellet was solubilized in 10 volumes of 7 M urea, 2 M thiourea, 4 % CHAPS, 30 mM Tris, pH 8.5 and cleared by centrifugation at 12000 x g, 4 °C for 45 mins. Protein concentrations of each myofibrillar and soluble protein sample were measured using the Bradford assay (Sigma-Aldrich, Poole, Dorset, United Kingdom). The protein concentrations of experimental samples were interpolated from bovine serum albumin standards (0.125 mg/ml – 1.0 mg/ml range) by linear regression. The total protein content (mg) of each muscle was calculated by multiplying muscle wet weight (WW; mg) by the yield of protein extracted from an aliquot of muscle powder (MP; mg) divided by the protein concentration (PC; mg/ml) and homogenate volume (HV; ml) of the myofibrillar and soluble protein fractions (Equation 1).

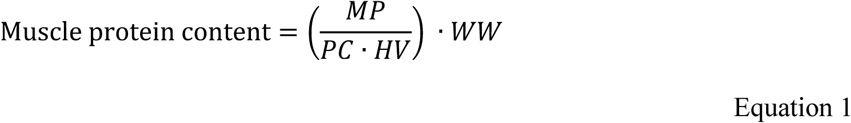

Muscle proteins were processed for mass spectrometry by tryptic digestion according to the filter aided sample preparation (FASP) method (Wiśniewski et al., 2009). Lysates containing 100 µg protein were precipitated in 5 volumes of acetone at -20 °C for 1 h. After centrifugation (5,000 x g, 5 min), acetone was decanted and the protein pellets were air dried and then resuspended in 200 µl of UA buffer (8 M urea, 100 mM tris, pH 8.5). Samples were incubated at 37 °C for 15 min in UA buffer with 100 mM dithiothreitol (DTT) followed by 20 min at 4 °C in UA buffer containing 50 mM iodoacetamide (protected from light). Samples were washed twice with 100 µl UA buffer and transferred to 50 mM ammonium hydrogen bicarbonate (Ambic). Sequencing grade trypsin (Promega; Madison, WI, USA) in 50 mM Ambic was added at an enzyme to protein ratio of 1:50 and the samples were digested overnight at 37 °C. To terminate digestion, peptides were collected in 50 mM Ambic and trifluoracetic acid (TFA) was added to a final concentration of 0.2 % (v/v). Aliquots containing 4 µg peptides, were desalted using C_18_ Zip-tips (Millipore, Billercia, MA, USA) and eluted in 50:50 of acetonitrile and 0.1 % TFA. Peptide solutions were dried by vacuum centrifugation for 25 min at 60 °C and peptides were resuspended in 0.1 % formic acid (FA) spiked with 10 fmol/µl yeast ADH1 (Waters Corp.) in preparation for LC-MS/MS analysis (Silva et al., 2006).

### Liquid chromatography-tandem mass spectrometry

Peptide mixtures were analysed using an Ultimate 3000 RSLC nano liquid chromatography system (Thermo Scientific) coupled to Q-Exactive orbitrap mass spectrometer (Thermo Scientific). Samples were loaded, via ulPickUp injection, on to the trapping column (Thermo Scientific, PepMap100, 5 μm C_18_, 300 μm X 5 mm) in 0.1 % (v/v) TFA and 2 % (v/v) ACN at a flow rate of 25 μl/min for 1 minute. Samples were resolved on a 500 mm analytical column (Easy-Spray C_18_ 75 μm, 2 μm column) and peptides eluted using a linear gradient from 97.5 % A (0.1 % formic acid) 2.5 % B (79.9 % ACN, 20 % water, 0.1 % formic acid) to 50 % A: 50 % B over 150 min at a flow rate of 300 nl/min. Data-dependent selection of the top-10 precursors selected from a mass range of m/z 300-1600 was used for data acquisition, which consisted of a 70,000-resolution (at m/z 200) full-scan MS scan (AGC set to 3e^06^ ions with a maximum fill time of 240 ms). MS/MS data were acquired using quadrupole ion selection with a 3.0 m/z window, HCD fragmentation with a normalized collision energy of 30 and in the orbitrap analyser at 17,500-resolution at m/z 200 (AGC target 5e4 ion with a maximum fill time of 80 ms). To avoid repeated selection of the same peptides for MS/MS, a dynamic exclusion window of 30 s was applied.

### Label-Free quantification of protein abundances

Progenesis Quantitative Informatics for Proteomics (QI-P; Nonlinear Dynamics, Waters Corp., Newcastle, UK) was used for label-free quantitation (LFQ), consistent with previous studies (Camera et al., 2017, Hesketh et al., 2020, Srisawat et al., 2023). MS data were normalized by inter-sample abundance ratio, and relative protein abundances were calculated using nonconflicting peptides only. MS/MS spectra were exported in Mascot generic format and searched against the Swiss-Prot database (2022_08) restricted to ‘Rattus’ (8,071 sequences) using a locally implemented Mascot server (v.2.8 www.matrixscience.com) with automatic target-decoy search. The enzyme specificity was trypsin with 2 allowed missed cleavages, carbamidomethylation of cysteine (fixed modification) and oxidation of methionine (variable modification). M/Z error tolerances of 10 ppm for peptide ions and 20 ppm for fragment ion spectra were used. The Mascot output (xml format), restricted to non-homologous protein identifications was recombined with MS profile data in Progenesis. Protein abundance data was calculated using only unique peptides with high confidence (<1 % false-discovery rate; FDR) identification scores. The mass spectrometry proteomics data have been deposited to the ProteomeXchange Consortium via the PRIDE partner repository with the dataset identifier PXD060879 and 10.6019/PXD060879.

### Measurement of body water D2O enrichment

Body water enrichment of D_2_O was measured in plasma samples against external standards constructed by adding D_2_O to PBS over the range from 0.0 to 5.0 % in 0.5 % increments. D_2_O enrichment of aqueous solutions was determined by gas chromatography-mass spectrometry after exchange with acetone (McCabe et al., 2006). Samples were centrifuged at 12,000 g, 4 °C for 10 min, and 20 µl of plasma supernatant or standard was reacted overnight at room temperature with 2 µl of 10 M NaOH and 4 µl of 5 % (v/v) acetone in acetonitrile. Acetone was extracted into 500 µl chloroform, and water was captured using 0.5 g Na_2_SO_4_. A 200 µl aliquot of chloroform was transferred to an auto-sampler vial, and samples and standards were analysed in triplicate using an Agilent 5973 N mass selective detector coupled to an Agilent 6890 gas chromatography system (Agilent Technologies, Santa Clara, CA, USA). A CD624-GC column (30 m x 0.53 mm x 3 μm) was used in all analyses. Samples (1 μl) were injected using an Agilent 7683 auto sampler. The temperature program began at 50 °C, increased by 30 °C/min to 150 °C and was held for 1 min. The split ratio was 50:1 with a helium flow of 1.5 ml/min. Acetone eluted at ∼3 min. The mass spectrometer was operated in the electron impact mode (70 eV), and selective ion monitoring of m/z 58 and 59 was performed using a 10 ms/ion dwell time. Percent D_2_O enrichment of experimental samples was calculated by interpolation from the standard curve using linear regression.

### Absolute dynamic profiling technique for proteomics (Proteo-ADPT)

The absolute dynamic profiling technique for proteomics (Proteo-ADPT) was conducted by calculating absolute protein abundance and synthesis rates on a protein-by-protein basis, originally described in Hesketh et al.,(Hesketh et al., 2020). Relative protein abundances (fmol/ μg protein on column) measured by LFQ were converted to absolute (μg) values using Equation 2.

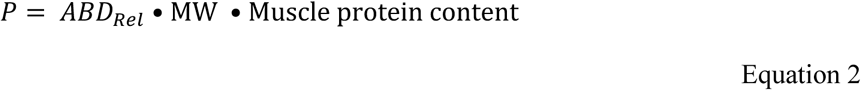

That is the absolute abundance (P; μg/ muscle) of each protein was calculated from the relative abundance (ABD_rel_; fmol/ μg total protein) measured by LFQ multiplied by the molecular weight (MW; kDa) of the protein (referenced from the UniProt Knowledgebase) and the total protein content (mg) of the muscle (derived in Equation 1).

Mass isotopomer abundance data were extracted from MS spectra using Progenesis Quantitative Informatics (Non-Linear Dynamics, Newcastle, UK). Consistent with previous work (Camera et al., 2017, Hesketh et al., 2020, Nishimura et al., 2023), the abundances of peptide mass isotopomers were collected over the entire chromatographic peak for each proteotypic peptide that was used for label-free quantitation of protein abundances. Mass isotopomer information was processed using in-house scripts written in Python (version 3.12.4). The incorporation of deuterium into newly synthesised protein was assessed by measuring the increase in the relative isotopomer abundance (RIA) of the m_1_ mass isotopomer relative to the sum of the m_0_ and m_1_ mass isotopomers (Equation 3) that exhibits rise-to-plateau kinetics of an exponential regression (Sadygov, 2020) as a consequence of biosynthetic labelling of proteins *in vivo*.

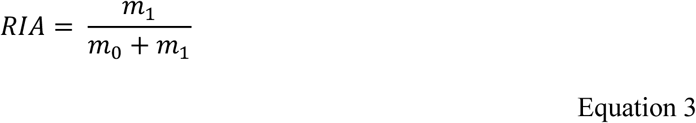

The plateau in RIA (*RIA_plateau_*) of each peptide was derived (Equation 4) from the total number (*N*) of ^2^H exchangeable H—C bonds in each peptide, which was referenced from standard tables (Holmes et al., 2015), and the difference in the D:H ratio (^2^H/^1^H) between the natural environment (*DH_nat_*) and the experimental environment (*DH_exp_*) based on the molar percent enrichment of deuterium in the precursor pool, according to (Ilchenko et al., 2019).

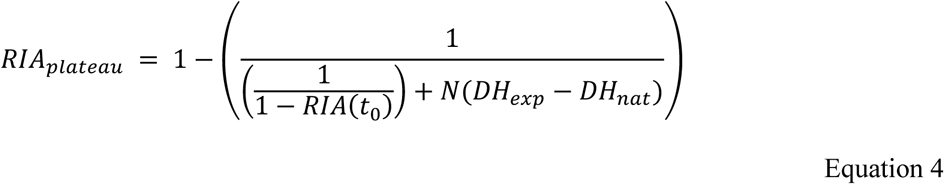

The rate constant of protein degradation (*k_deg_)* was calculated (Equation 5) between the beginning (t_0_) and end (t_1_) of each 10-day labelling period. Calculations for exponential regression (rise-to-plateau) kinetics reported previously (Ilchenko et al., 2019) were used and *k_deg_* data were adjusted for differences in protein abundance (P) between the beginning (t_0_) and end (t_1_) of each labelling period.

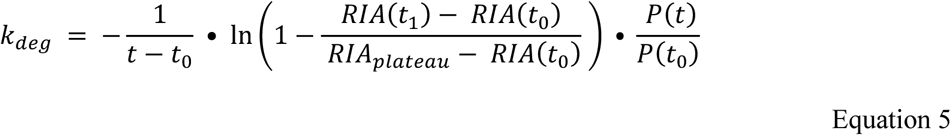

Absolute synthesis rates (ASR) were derived (Equation 6) by multiplying peptide *K_deg_* by the absolute abundance (e.g. μg protein/ muscle) of the protein at the end of the labelling period P(t).

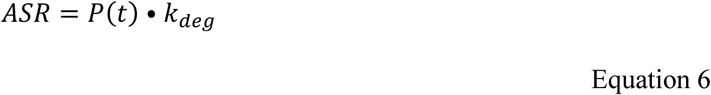

Absolute degradation rates (ADR) were derived (Equation 7) by subtracting the rate of change in abundance from the absolute synthesis rate.

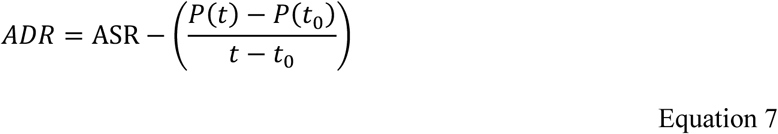

### Statistical and bioinformatic analysis

Unless stated otherwise, data are presented as mean ± standard deviation (SD) and statistical analyses were conducted in *R* (version 4.3.1). Muscle wet weight and protein content were normalised to animal body weight and analysed using mixed analysis of variance (ANOVA). Where appropriate (i.e. non-omics data) significant (p <0.05) interactions were further investigated by post-hoc pairwise comparisons of estimated marginal means with Bonferroni adjustment.

Proteomic data were filtered to exclude proteins that were not quantified in all conditions (Stim and Ctrl) at all experimental time-points (0, 10, 20, and 30 d) in all biological replicates (n = 4, in each group), i.e. proteins with missing values were removed and no data were imputed. Two-factor mixed ANOVA was used to identify significant interactions between condition (PRT stimulated vs contralateral control) and time (experimental time points; 0 d, 10 d, 20 d and 30 d). In addition, within-subject ANOVA was used to investigate differences between PRT stimulated and contralateral control muscle at each experimental timepoint or period. Type I error was controlled using the calculation of false-discovery rates (q values) and BH-corrected P values using the ‘qvalue’ package (R/Bioconductor) (Storey and Tibshirani, 2003).

Relationships between changes in protein abundances and/ or absolute synthesis rates were assessed using linear regression at single experimental time points or multiple linear regression across multiple experimental time points. Regression models predicting changes in absolute protein abundance from absolute synthesis rate data were constructed from log_2_-transformed fold-difference (Stim/ Ctrl) at each experimental time point.

Proteins that exhibited statically significant (BH-corrected P <0.05) interactions (condition x time) in abundance were submitted to bioinformatic analyses to further explore differences in the temporal profile of protein abundance changes. To enable exploration of temporal profiles of individual protein synthesis rates, unsupervised clustering was conducted across all protein-specific ASR data to identify patterns and networks of proteins with similar synthesis rate dynamics across the 30-day experimental period. Temporal profiles (log_2_ fold-difference Stim/Ctrl at days 0, 10, 20, and 30) were normalised using the ‘standardise()’ function within the mFUZZ R package (Kumar and Futschik, 2007) and submitted to soft clustering analysis using the fuzzy c-means clustering algorithm. The optimal number of clusters was determined by inspection of scree plots and qualitative iterative assessment of cluster membership profiles. The minimum membership value for inclusion into a cluster was set at 0.5 throughout. Proteins exhibiting significant differences between Stim and Ctrl across the timecourse and subsequent temporal clusters were further investigated using bibliometric mining in the Search Tool for the Retrieval of INteracting Genes/proteins (STRING, Version 12) using the evidence of interaction sources of bibliometric textmining, experimental verified protein-protein interaction data, gene ontology databases, and co-expression data with the minimum required interaction score set at 0.4 (medium confidence). Protein-Protein interaction networks and functional enrichment analyses were conducted using the STRINGdb package in R (version 2.12.1) (Szklarczyk et al., 2019) and corrected against the experiment-specific background consisting of all proteins that were included in statistical analysis. Protein-protein interaction networks were transferred via the RCy4 R packaged (Version 1) and visualised using Cytoscape version 3.9.1 (Shannon et al., 2003).

## Results

### Daily PRT increases TA wet weight and protein content

TA wet weight and protein content each showed a significant (p <0.001) interaction between condition (Stim vs Ctrl) and time during the 30-day PRT intervention. Post-hoc analyses confirmed there was no difference in either TA wet weight or protein content between the left (sham-operated) and right (contra-lateral control) limbs at the onset (Day 0) of the experiment. Compared to Ctrl TA, significant (p <0.05) increases in wet weight (Figure 1B) and total protein content (Figure 1C) were evident in the Stim TA after 10 days PRT (+15.52 ± 7.79 % and +30.48 ± 8.56 %, respectively) and these differences were sustained across the 20 d (+19.48 ± 3.22 % and +50.21 ± 12.68 %, respectively) and 30 d time points (+18.13 ± 7.93 % and +40.02 ± 10.99 %, respectively). Changes in muscle wet weight were significantly (p = 1.156e^-05^) correlated with changes in muscle protein accretion (Pearson correlation coefficient (*r*) = 0.87, 95 % Confidence interval = 0.66 – 0.95) (Figure 1D). Changes in muscle wet weight were therefore due to increases in protein content (growth) rather than oedema or increase in water content that can accompany a response to muscle damage.

**Figure 1.**
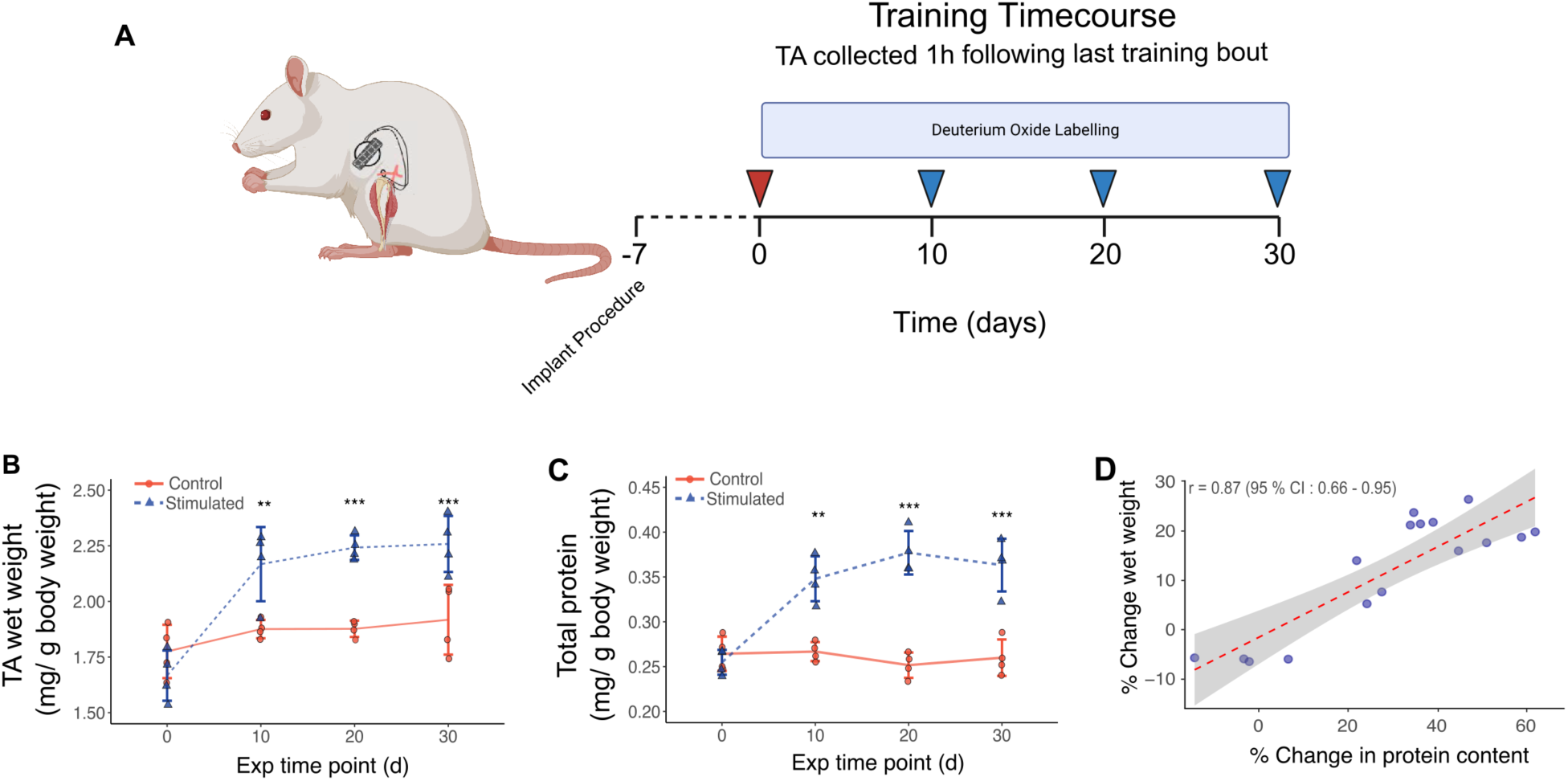
Daily programmed resistance training increases TA wet weight and protein content. (**A**) Programmed resistance training (PRT) *in vivo* was achieved using implantable pulse generators placed in the abdomen with electrodes routed subcutaneously to the left limb. Within-animal unilateral PRT model (n = 4 per time point), uses loaded contractions of the tibialis anterior (resisted by the plantarflexors) in the left hindlimb whilst the right remains unstimulated. Rats were stimulated daily during the early phase of the light period. Control and sham-operated control TA muscles were collected (red arrow) prior to the initiation of deuterium oxide labelling on day 0. Deuterium oxide (D_2_O) labelling was initiated at day 0 and maintained by administration of D_2_O in the drinking water available ad libitum. Stimulated and control limb muscles were collected 1 h after the last training bout following 10, 20, or 30 days of daily training. PRT consisted of 5 sets of 10 repetitions of 2 s on, 2 s off, tetanic contractions at 100 Hz with 2.5 min of rest between sets. (B) Time course changes in TA mass (TA wet weight relative to body weight; mg/ g) of the stimulated and control limbs across the experimental time-course **(C**) and protein content (relative to body weight; mg/ g) measured by Bradford assay. Data are presented as Mean ± Standard deviation and analysed using a two-way ANOVA. Significant interactions (Condition x Time) were investigated using pairwise comparisons with Bonferroni adjustment * P ≤ 0.05, ** P ≤ 0.01, *** P ≤ 0.001. (**D**) Correlation analysis of changes in TA wet weight and protein content. Data points represent mean percentage change (Stim/ Ctrl) from n = 4 biological replicates across 0, 10, 20, and 30 days.

### The proteome of contralateral (non-stimulated) control muscle was unaffected by PRT

Proteomic analysis confidently identified and quantified the abundance of 1083 proteins in rat TA. Prior to statistical analyses, protein abundance data were stringently filtered to exclude proteins that were not quantified in all Stim and Ctrl muscles (n = 4, in each group) across all experimental time points (days 0, 10, 20, and 30). After filtering, 658 proteins were quantified in all 32 biological samples. Protein abundances (expressed as mass of protein in each TA) spanned from 0.49 ± 0.32 ng (Histone H1.1) to 28.514 ± 3.846 mg (Myosin heavy chain 4) and the top 10 most abundant proteins accounted for >50 % (∼53 mg) of total protein content (92 mg) in Ctrl muscles collected on day 0 (Figure 2A).

**Figure 2.**
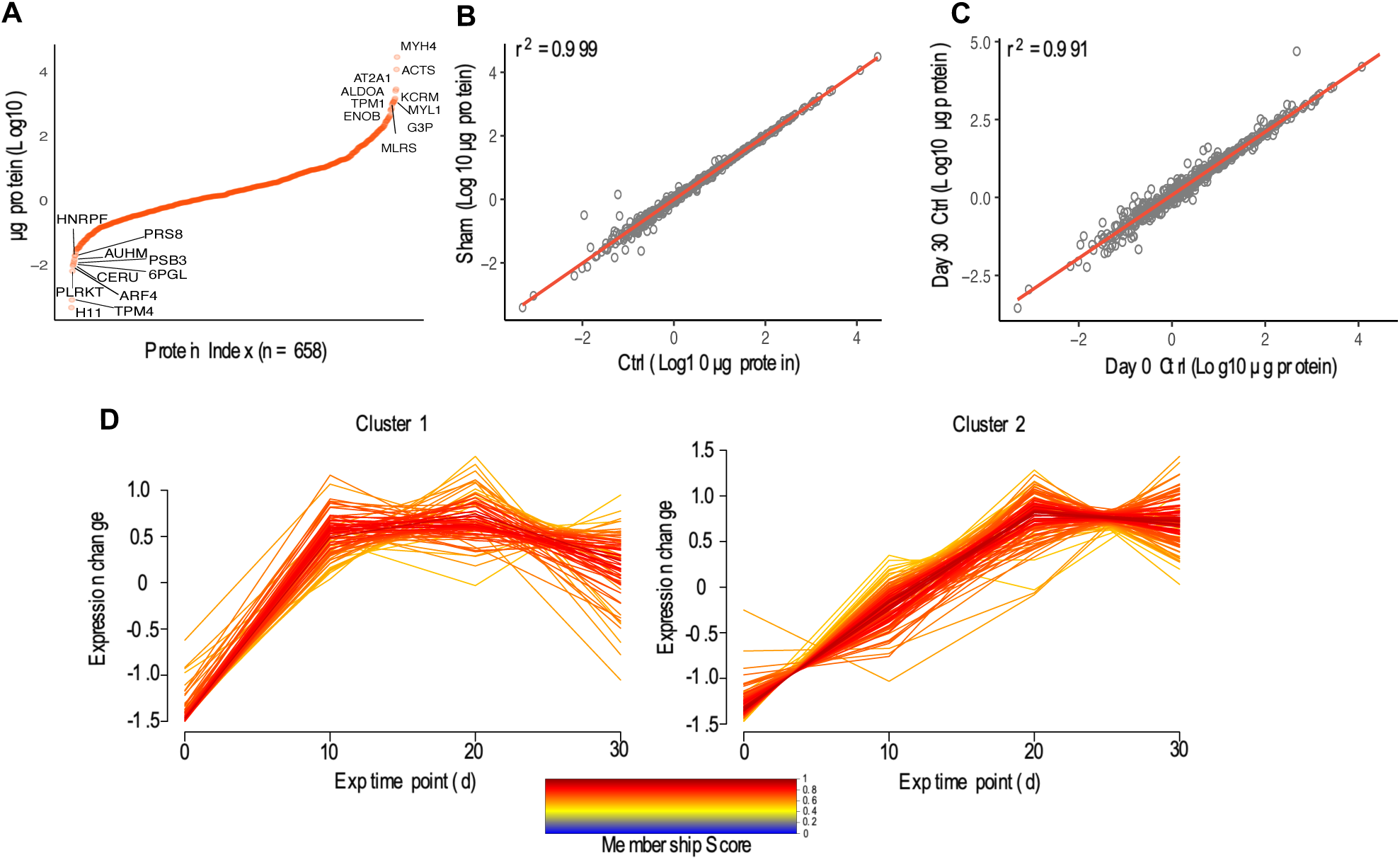
Absolute quantification of muscle proteins in rat TA. **(A)** Distribution plot of 658 proteins successfully quantified (µg of protein) across all 32 muscle samples. Data points represent mean µg abundance from Ctrl TA at the onset of the experiment (n = 4). (**B**) Ordinary least squares (OLS) regression analysis (r^2^) of absolute abundance (µg) of n = 658 proteins quantified in TA of n =4 sham-operated and contralateral Ctrl limbs at the beginning (day 0) of the experimental period. (**C**) OLS (r^2^) in protein abundances (n = 658) quantified in Ctrl TA at the beginning (day 0) and end (day 30) of the experimental period (n = 4, biological replicates). (**D**) Soft clustering analysis displays temporal profiles (log_2_ transformed Stim/Ctrl fold-difference at each time point) of proteins exhibiting significant interactions (BH adj P < 0.05) in abundance, using the fuzzy c-means clustering algorithm with the Mfuzz R package. The minimum membership value for inclusion into a cluster was set at 0.5.

At the onset of the experiment there were no significant differences in abundance for any protein between the sham-operated and contralateral Ctrl muscle. The coefficient of determination (*r^2^*) for protein abundance between the Sham operated and Ctrl muscle on day 0 was *r^2^* = 0.999 (Figure 2B). The median (M) coefficient of variation (CV) of absolute protein abundances quantified between Sham operated and Ctrl TA at the onset of the experiment was 4.6 % (1^st^ quartile = 2.04 % and 3^rd^ quartile = 6.64 %). Ordinary Least Squares regression indicated a strong linear relationship between the absolute abundances quantified in Ctrl TA at onset of the experiment (day 0) and day 10 (r^2^ = 0.995), day 20 (r^2^ = 0.989), and day 30 (r^2^ = 0.991) (Figure 2C). We are confident this demonstrates a high level of consistency across the independent groups of Ctrl muscle across the 30-d experimental period, and that the contralateral (unstimulated Ctrl) muscle in the stimulated animals is little effected by daily stimulation in the contralateral limb.

### Ribosomal proteins respond early to PRT

Two-way ANOVA (Time*Condition) indicated robust (BH-corrected P < 0.05) changes in the abundance profiles of 187 proteins in Stim muscle across the 30-d training period. Unsupervised soft-clustering analysis of the 187 significant proteins highlighted 2 prominent temporal patterns amongst the proteins that responded to PRT (Figure 2D).

Proteins in Abundance Cluster 1 (ABD_Cluster_1; 74 in total) exhibited increases in absolute abundance after the first 10 days of PRT and then plateaued in abundance after 20 days or 30 days PRT (Figure 2D). Gene Ontology Biological Processes including post-transcriptional regulation and translation of mRNA (Figure 3A and 3B) were significantly enriched amongst proteins in ABD_Cluster_1, which included 18 ribosomal subunits encompassing 10 large (60S) and 8 small (40S) ribosomal subunits. Eukaryotic translation initiation factors (eIF), eIF3A and eIF2G were also included within ABD_Cluster_1 alongside the eukaryotic translation elongation factor (eEF) eEF1D and the peptide chain release factor, ERF1. In addition, ABD_Cluster_1 contained a network of proteins associated with skeletal muscle development and organisation including 4 tubulin proteins (TBA1B , TBA4A, TBB4A, and TBB5), dynactin subunit 1 and 2, Nexilin, WD repeat-containing protein 1, Alpha-crystallin B chain, Vimentin, Prelamin-A/C, Tropomodulin-1, and 5 non-typical myosin isoforms (MY18A, MYH11, MYH8, MYH9, and MYO1C).

**Figure 3.**
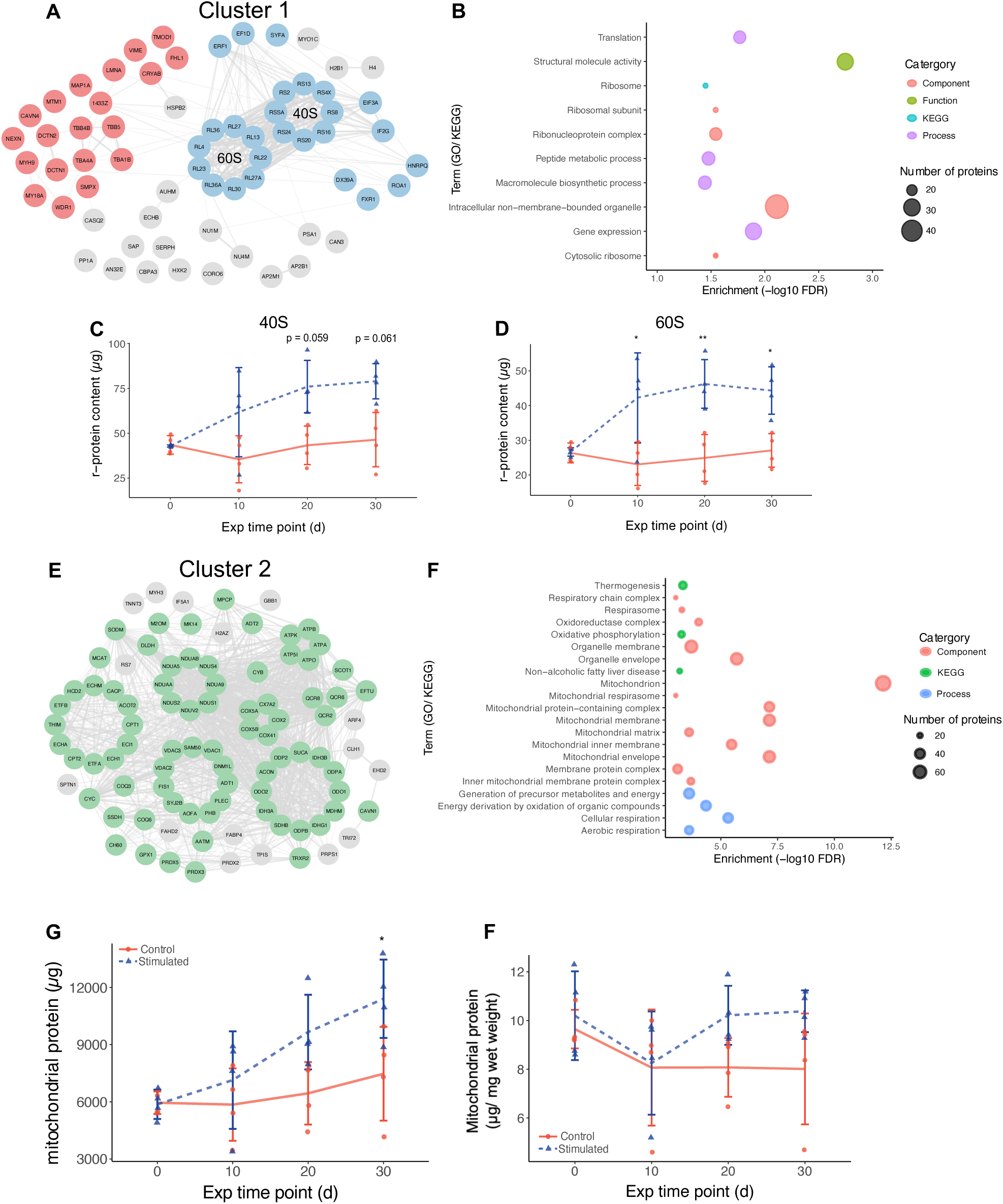
Ribosomal protein accretion occurs before mitochondrial proteome responses. **(A)** STRING protein interaction network for proteins displaying significant (BH-adj P < 0.05) interaction (Time*Condition) in abundance data across the time-course of PRT belonging to cluster 1 reported in Figure 2. Coloured nodes represent proteins in cluster 1 associated with sarcomere organisation and muscle contractions (Red) and protein translation (Blue). (**B**) Bubble plot represents significantly enriched (FDR < 0.05) Gene ontology (GO) and KEGG pathway terms for proteins included in cluster 1. Time course changes in total ribosomal protein content (µg) of the ‘small’ 40S (**C**) and ‘large’ 60S (**D**) subunits. (**E**) STRING protein interaction network for proteins displaying significant (BH-adj P < 0.05) interaction (Time x Condition) in abundance data across the time-course of PRT belonging to cluster 2 reported in Figure 2. Coloured nodes represent proteins in cluster 2 associated with the GO cellular component ‘Mitochondrion’ (Green). (**F**) Bubble plot represents significantly enriched (FDR < 0.05) Gene ontology (GO) and KEGG pathway terms for proteins included in cluster 2. **(G)** Time course changes in total mitochondrial protein content expressed in absolute units (µg protein) and **(H)** relative to TA wet weight (µg protein/ mg) (**H**). Data in **C**, **D**, **F**, and **H** are presented as Mean ± Standard deviation and analysed using a two-way ANOVA. Significant interactions (Condition x Time) were investigated using pairwise comparisons with Bonferroni adjustment. * P ≤ 0.05, ** P ≤ 0.01, *** P ≤ 0.001.

In all, our analysis quantified the abundance of 57 out of the 80 annotated subunits of the ribosome, including 30 subunits of the 60S large ribosome and 27 subunits of the 40S small ribosome. Changes in the sum abundance of ribosomal proteins (r-protein) in response to PRT mirrored the accretion of total muscle protein (Figure 3C and 3D) and changes in the total content of the 40S and 60S subunits exhibited a significant (P = 0.047 and 0.0097, respectively) interaction between condition (PRT vs Ctrl) and experimental time point (day 0 vs day 30). Post-hoc analysis indicated that after 10 or more days PRT, the total r-protein content of the 60S subunit was 84 % greater (p = 0.01) compared to the contralateral Ctrl, whereas the total abundance of 40S subunits tended (p = 0.06) to be 76 % greater compared to Ctrl after 20 days PRT (Figure 3C and 3D).

### Mitochondrial protein response occurs after ribosomal protein response

Proteins in ABD_Cluster_2 exhibited more gradual rises (compared to ABD_Cluster_1) in abundance and did not exhibit statistically relevant differences between Stim and Ctrl condition until after 20 days and 30 days PRT (Figure 2D). ABD_Cluster_2 included 114 mitochondrial proteins (from 162 proteins assigned to Cluster 2) and was significantly enriched (FDR < 0.001) in a collection of GO terms associated with mitochondria and cellular respiration (Figure 3E and 3F) including 23 proteins associated with oxidative phosphorylation, comprising 9 respiratory Complex I subunits and 6 ATP synthase subunits. The GO Biological Processes ‘Mitochondrion Organization’ was significantly enriched (FDR = 0.01, 23 proteins) within ABD_Cluster_2 and included key regulators of mitochondrial quality control (e.g. VDAC-1, -2, and -3, FIS1, SAM50, SODM, PHB, and CH60). Cluster 2 was also significantly enriched with proteins associated with the KEGG pathway TCA Cycle (FDR = 0.01, 13 proteins), and ABD_Cluster_2 contained eIF5A1 and the mitochondrial elongation factor TU (Figure 3E).

Total mitochondrial protein content was estimated by aggregating the absolute abundance (µg of protein) of all proteins annotated to the GO Cellular Component ‘Mitochondrion’ within our analysis. Two-way ANOVA identified a significant (p = 0.041) interaction (Time*Condition) in total mitochondrial protein content (Figure 3G). Post-hoc analysis revealed significant (p = 0.037) increases in total mitochondrial protein content between the Stim and Ctrl TA were only evident after 30 days PRT.

### Absolute quantification of proteome dynamics

Proteo-ADPT was conducted on a subset of proteins that had high-quality peptide mass isotopomer profiles. In total we quantified the rates of synthesis for 585 proteins. After filtering for missing values, the absolute and fractional synthesis rate (ASR; ng/d and FSR; %/d, respectively) of 215 proteins were quantified in all samples (n = 4 Ctrl and Stim replicates) across the early (day 0 – day 10), intermediate (day 10 – day 20) and later (day 20 – day 30) measurement periods. Proteins included in the Proteo-ADPT analysis were among the most abundant in TA muscle and shared gene ontology terms including the biological processes ‘Metabolic Process’ (159 proteins), ‘Cellular Respiration’ (46 proteins), cellular components ‘Mitochondrion’ (89 proteins) and ‘Myofibril’ (35 proteins). In sum, the 215 proteins included in the Proteo-ADPT analysis represented ∼70 % of total muscle protein content. In Ctrl muscles, the total amount of new protein synthesised was 2.61 ± 1.63 mg/d throughout the experimental period, which equates to an average mixed-protein FSR of 4.26 ± 2.17 %/d. Two-way mixed ANOVA did not identify significant (NS; P>0.14) effects of condition (Stim vs Ctrl) or interactions between condition and measurement period (0-10 d, 10-20 d or 20-30 d) in either total ASR or mixed-protein FSR. Mixed-protein FSR was ∼1.63 fold greater in Stim (5.17 ± 1.48 %/d) compared to Ctrl TA (2.91 ± 0.68 %/d) during the early (0-10 d period) response of PRT (Figure 4A and B), which equates to a non-significant rise of ∼1.80 mg/d total protein synthesised (total ASR) in the Stim (3.53 ± 2.12 mg/d) compared to the Ctrl TA (1.73 ± 0.54 mg/d). Translational efficiency (µg protein synthesised per µg r-protein content; µg ASR/ µg r-protein) was assessed by comparing the total ASR (mg/ d of protein synthesis) relative to r-protein content in each muscle and did not change in response to PRT. Translational capacity (µg ASR/ µg r-protein) in exercised (Stim) muscle during the early (31.53 ± 10), mid (39.38 ± 17.75) or late (19.55 ± 5.87) experimental periods was not different from values in Ctrl muscle (Figure 4C) during the early (29.71± 1.37), mid (45.04 ± 28.09) or late (38.09 ± 16.10) experimental periods.

**Figure 4.**
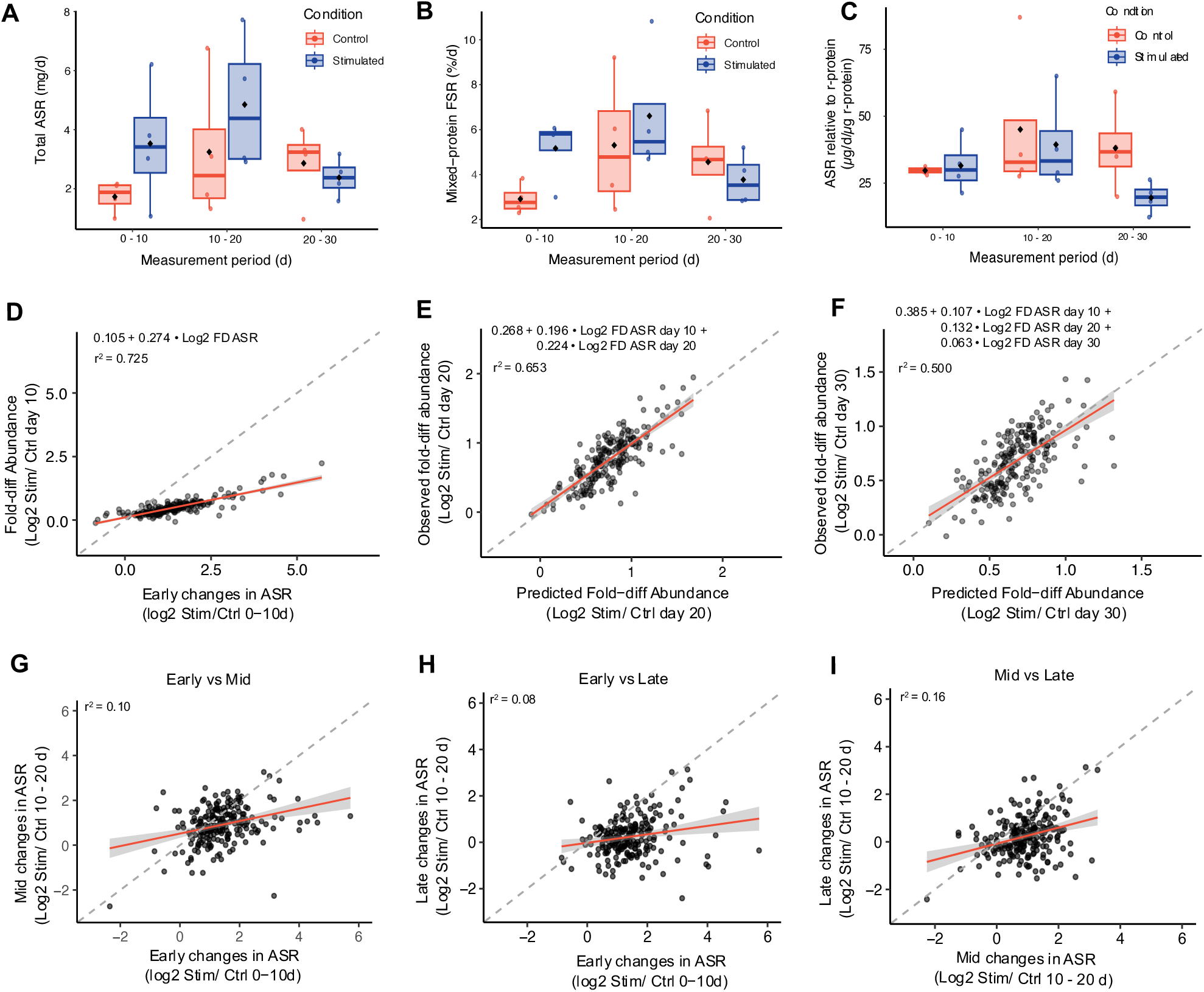
Absolute quantification of protein synthesis rates during PRT. (**A**) Boxplots of total muscle protein synthesis rates (sum of ASR) quantified from 216 protein-specific synthesis rates quantified across the entire experimental design. (**B**) Mixed-protein synthesis rates represented as fractional synthesis rates (FSR; %/d) calculated from median FSR of 216 proteins quantified across the entire experimental design. (**C**) Estimates of translational efficiency in Stim and Ctrl muscles across the time course of PRT expressed as total muscle protein synthesis relative to total ribosomal protein content (µg/ d/ µg of r-protein). Data analysed using a two-way ANOVA. Black diamond represents mean value. (**D**) Ordinary least squares regression analysis (OLS) of log^2^ fold-difference in ASR (Stim/ Ctrl) across the early (0 – 10d) period of PRT and the measured log^2^ fold-difference in abundance (Stim/ Ctrl) on day 10 (n = 216 proteins). Multiple regression analysis of changes in protein abundance (log^2^ Stim/ Ctrl) after 20 (**E**) and 30 days (**F**) of PRT. Predicted measurement changes (log^2^ Stim/ Ctrl) in protein specific ASR across early and mid (**E**) and early, mid, and late experimental periods. **(G-I**) OLS of changes in ASR (log^2^ Stim/ Ctrl) for 216 individual proteins measured across the entire experimental period in Stim and Ctrl TA.

Protein production (i.e. ASR) outweighed protein accretion during the early (day 0 – day 10) adaptive response to PRT (Figure 4D). There was a strong relationship (r^2^ = 0.725) between the difference (within animal Stim/ Ctrl) in ASR and change in protein abundance, but the slope (0.274) of the linear regression suggests only 27% of the protein synthesised was accreted into muscle protein. Multiple linear regression on changes in ASR across both the early- and mid- (Figure 4E) or during early-, mid- and late- (Figure 4F) experimental periods explained ∼65 % of the gain in muscle protein content after 20 days PRT. Whereas the elevation in ASR across early-, mid-, and late-experimental periods explained ∼50 % of the gain in protein content after 30 days of PRT (Figure 4F). The linear relationship (log_2_ fold-difference abundance = 0.274 • log_2_ fold-difference ASR + 0.105) between changes in ASR and abundance across the first 10 days indicates ‘over-synthesis’ of proteins during muscle growth, i.e. during the first 10 days PRT a log_2_ fold-increase in ASR of 1 equated to log_2_ fold-increase in abundance of ∼0.3.

Unsupervised clustering and network analyses of changes in individual protein ASR (Figure 5A-B) highlighted 3 patterns of response. ASR_Cluster _1 comprised 44 proteins that are upregulated in ASR specifically during the early (0-10 d) response to PRT, including proteins involved in translation (6 ribosomal proteins: RS-SA, - 6, -9, -19, -20, and RL6 and eEF1G, eF2, and EFTU). ASR_Cluster_1 also contained proteins associated with muscle growth and development (MYH3, MYH8, CRYAB, TRDN and DESM (Figure 5B, and glycolytic enzymes (Fructose-bisphosphate aldolase A; ALDOA, B-enolase; ENOB, Pyruvate dehydrogenase E1 component subunit alpha; ODPA, Dihydrolipoyllysine-residue acetyltransferase component of pyruvate dehydrogenase complex; ODP2, Phosphorylase b kinase gamma catalytic chain; PHKG2, Glycogen Phosphorylase; PYGM, Phosphoglucomutase-1; PGM1, and ATP-dependent 6-phosphofructokinase, muscle type; PFKM).

**Figure 5.**
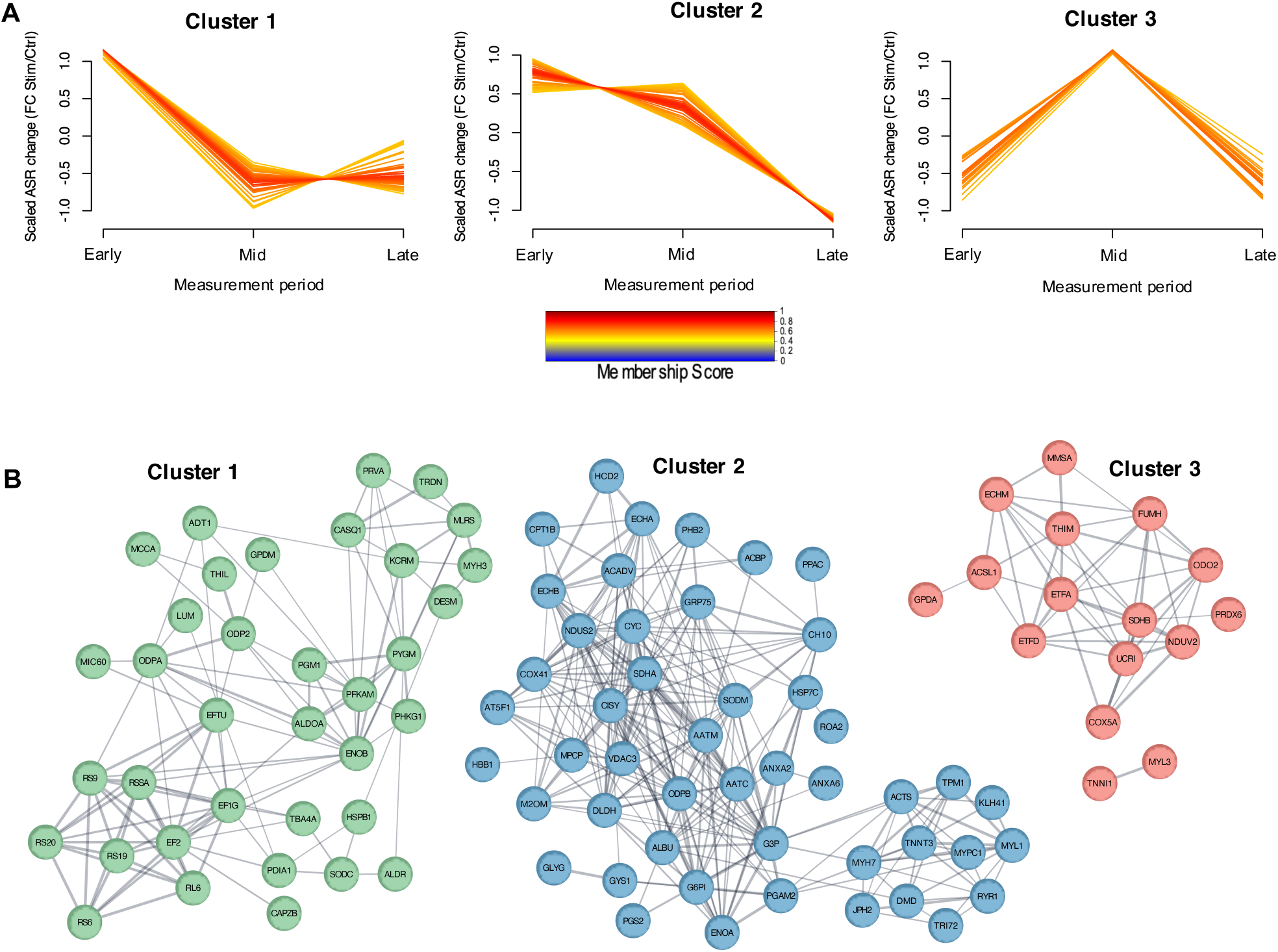
Time-course changes in synthesis rates occurring during PRT induced muscle growth. (**A**) Temporal profiles (log2 transformed Stim/Ctrl fold-difference at each time point) of individual protein ASRs underwent soft clustering analysis using the fuzzy c-means clustering algorithm with the Mfuzz R package. The minimum membership value for inclusion into a cluster was set at 0.5. (**B**) STRING protein interaction network for proteins contained in Cluster 1 (green), Cluster 2 (blue), and Cluster 3 (red) reported in panel B.

Proteins in ASR_Cluster_2 (54 proteins) exhibited greater ASR during both early- and mid-compared to the late (20-30 d) experimental periods and contain 2 subnetworks of proteins associated with muscle contraction (12 proteins including: DMD, TPM1, TNNT3, MYH7, ACTS1, MYL1, MYBPC1, AT2A1, JPH2, RYR1, KLH41 and CACB1) and a network of 24 proteins annotated to the GO cellular compartment ‘mitochondrion’ (Figure 5B. Proteins in ASR_Cluster_3 (19 proteins) exhibited greater ASR in exercised muscle specifically during the mid period and the majority (17/19 proteins) were associated with metabolic process, including 13 proteins annotated to the mitochondria (Figure 5B).

Twenty-seven proteins exhibited evidence of an interaction (p <0.1) between time (experimental period) and condition (Stim vs Ctrl) on changes in ASR, including 15 mitochondrial proteins (e.g. ATPA, ATPB, CPT1B, PRDX5, VDAC1, VDAC2, NDUA9, NDUAA) included in ABD_Cluster_2 as well as myofibrillar proteins (e.g. CRYAB and FLNC) included in ABD_Cluster_1. The ASR of ATP synthase subunit-β (ATPB; included in ABD_Cluster_2) exhibited a significant (p = 0.037) interaction (Time*Condition) and had a 3.6-fold greater rate of synthesis compared to Ctrl during the mid-period (10 d – 20 d) of PRT. In Ctrl muscle, ASR of ATPB was 8.40 ± 4.90 µg/d (between day 10 and day 20), the rate of change in ATPB abundance was 3.15 ± 7.76 µg/d, and the calculated degradation rate was 5.25 ± 2.91 µg/d. In Stim muscle, during the same d 10 – d 20 period, the ASR of ATPB was 30.39 ± 18.67 µg/d (approximately 22 µg/d greater than Ctrl), ATPB abundance increased at a rate of 17.35 ± 12.36 µg/d and the calculated degradation rate was 13.04 ± 6.97 (∼2.5-fold greater than Ctrl) (Figure 6A-C).

**Figure 6.**
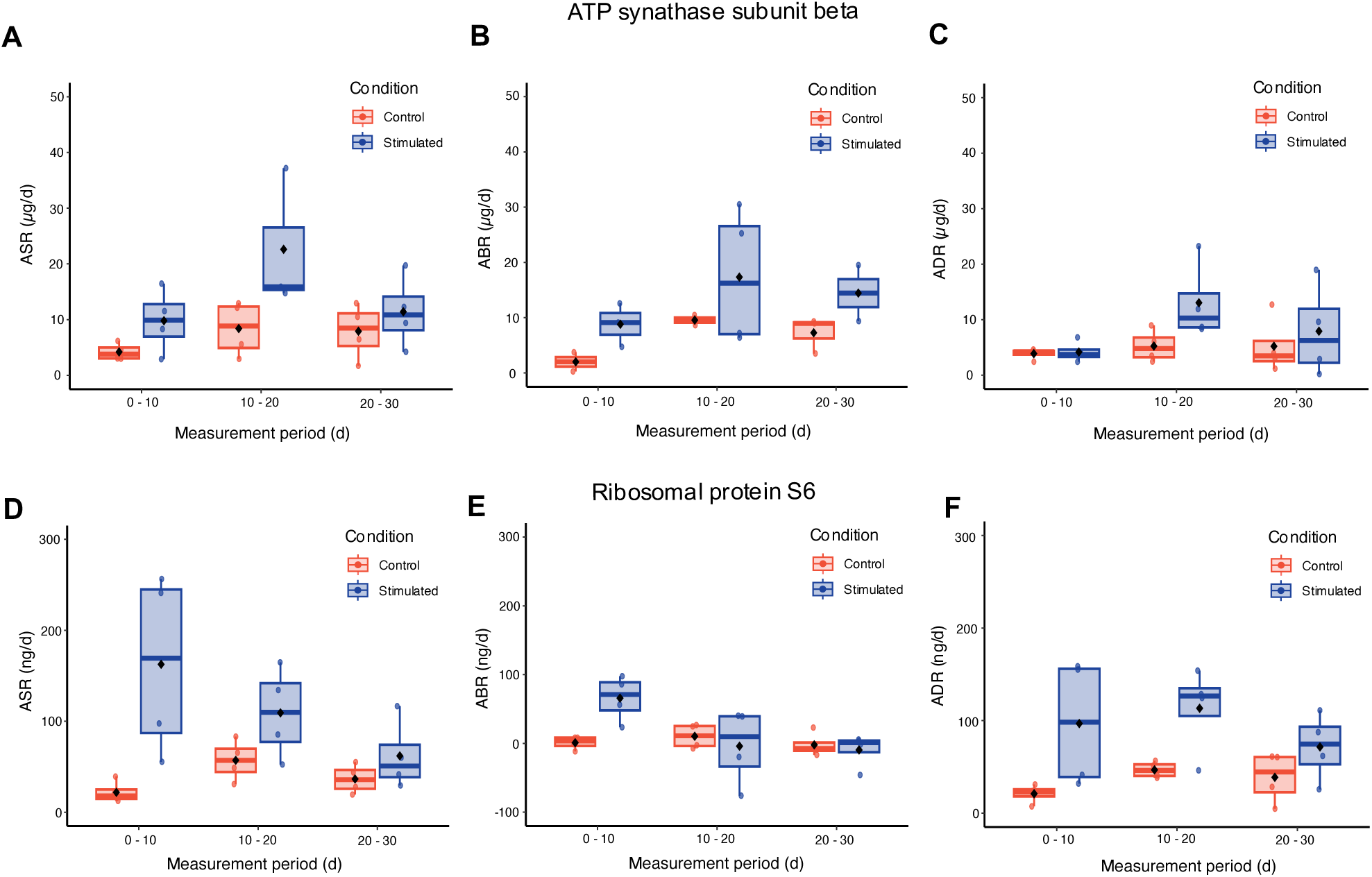
Proteo-ADPT analysis during PRT induced muscle growth. Boxplots of Proteo-ADPT data quantified across across the entire experimental design for ATP synthase subunit beta (**A-C**) and ribosomal protein S6 (**D-F**), including absolute protein synthesis rate (ASR; **A & D**), abundance rate of change (ABR; **B & E**), and absolute protein degradation rate (ADR; **C & F**). Data analysed using a two-way ANOVA. Black diamond represents mean value.

Ribosomal protein S6 (RS6) protein content accumulated at a rate of 65.77 ± 33.16 ng/d in Stim compared to 0.933 ± 9.47 µg/d in Ctrl across the first 10 days of PRT, in line with accumulation of ribosomal protein across the early adaptive period. The net increase in RS6 protein abundance was underpinned by a trend (P = 0.08) of Time x Condition interaction characterised by a 7.4-fold increase in ASR (Stim = 162.61 ± 101.01 and Ctrl = 21.96 ± 12.00 ng/d) and 4.6-fold increase in ADR (Stim = 96.83 ± 69.49 and Ctrl = 46.78 ± 8.69) during the first 10 days of PRT. This shows a concomitant increase in both the abundance and turnover rate of RS6 (Figure 6D-F).

## Discussion

Muscle proteome adaptations to resistance training (RT) are the foundation for improvements in muscle mass and strength, but our knowledge on the process of how the proteome adapts to RT remains incomplete. Gains in muscle mass are the prominent and desired outcome of RT and, in contrast to muscle responses to endurance training, RT is less associated with changes in muscle phenotype and proteome profile (Emanuelsson et al., 2024). Attempts to study muscle adaptations to RT using standard proteomic techniques, are constrained to only the relative abundance of muscle proteins and overlook co-occurring changes to the overall mass of the muscle. In addition to measurements on protein synthesis rates, Proteo-ADPT offers an opportunity to gain insight into the accretion of muscle proteins and reports changes to the absolute abundance profile of proteins occurring during the time course of adaptation to RT. Using unilateral programmed resistance training (PRT), we identified distinct temporal patterns of adaptation amongst different components of the skeletal muscle proteome. Unsupervised data analyses highlighted two distinct temporal patterns of adaptation during a 30-day experimental period. Networks enriched for translation- and ribosome-associated proteins were increased during the first 10 days of PRT (Figure 3A and 3B), whereas accretion of mitochondrial proteins occurred later in the adaptive response to PRT (Figure 3E-G).

Mechanical loading by RT enhances translational capacity in rat muscle as it does in human (Figueiredo et al., 2015), which likely underpin gains in muscle mass. Synergist ablation has been a key experimental model for studying muscle hypertrophy and protein accretion and, in rats, is associated with increases in translational capacity evidenced by increases in RNA/rRNA ratio (Nakada et al., 2016) and total RNA content of hypertrophied muscle (Roberts et al., 2020). Early work using electrically stimulated contractions in the lower limb of rats under anaesthesia also reported a 33 % increase in RNA concentration following 10 weeks of twice weekly electrical stimulation (Wong and Booth, 1990). Ribosomal proteins (r-proteins) have been less studied but have essential roles in rRNA processing, including ribosome maturation, assembly, stability and translational fidelity (Razi and Ortega, 2017). Our time-course clustering identified co-regulation of 18 r-proteins (8 small and 10 large subunit proteins) that significantly increased during the first 10 days of muscle growth. Similarly, the total protein content of the 40S (sum of 26 proteins) and 60S (sum of 30 proteins) ribosomal subunits exhibited comparable response patterns across the 30-day PRT period, indicating no significant alterations in r-protein stoichiometry during PRT-induced muscle growth.

Absolute data are impractical to acquire in humans, but increases in the relative abundance of ribosomal proteins have been reported after 4 weeks of thrice weekly resistance training (Jessen et al., 2024). Whereas no changes in ribosomal content were evident in proteomic analysis of muscle after a longer term, 8-week, RT intervention (Roberts et al., 2024). Herein, the early (0-10 days) response to PRT was dominated by increases in the abundance of proteins associated with translational capacity, which meant translational efficiency (i.e. total protein production per µg of r-protein) remained stable at ∼30 µg of protein synthesised per µg of r-protein per day. R-protein abundance is tightly controlled and gains in r-proteins plateaued after day 10 and remained stable across the final 20 days of PRT. Excess ribosomal subunits may be rapidly degraded by the ubiquitin proteasome system (Lam et al., 2007) but protein specific analysis of RS6 (Figure 6) suggests the plateau in ribosomal protein abundance was brought about by changes to both the synthesis and degradation rate. Compared to the early adaptive period, the RS6 protein ASR fell by 33 % in Stim TA and ADR increased by 17 % in the mid (10-20 days) experimental period, and both the ASR and ADR of RS6 further declined during the late (days 20-30) period.

Mitochondrial adaptations are universally accepted in response to endurance exercise, whereas the impact of RT on mitochondrial biogenesis and content are still debated (Parry et al., 2020). Early electron microscopy studies suggested mitochondrial volume may be diluted by gains in muscle mass associated with RT (MacDougall et al., 1979, Lüthi et al., 1986), whereas data on the relative abundance of mitochondrial proteins, such as citrate synthase, suggest there are no changes in mitochondrial volume in RT muscle (Groennebaek and Vissing, 2017). Gains in muscle mass induced by the ablation (Roberts et al., 2020) or tenotomy (Goldberg, 1968) or synergist muscles are associated with elevations in the fractional synthesis rate of protein mixtures from both the myofibrillar and soluble fractions of rat muscle, which may each include mitochondrial proteins. Our Proteo-ADPT analysis highlights mitochondrial aadaptation is a prominent component of the muscle response to daily PRT. Total mitochondrial protein content increased in line protein-specific responses in Abundance Cluster 2 (Figure 3G), which suggests reasonably coordinated gains in mitochondrial protein rather than stoichiometric remodelling of the mitochondrial proteome. ATP synthase subunits alpha (ATPA) and beta (ATPB), exhibited a >2-fold increase in synthesis in response to PRT during the intermediate (10-20 days) period whereas, in our (Viggars et al., 2023) transcriptomic analysis of this model, increases in mRNA expression of mitochondria- and oxidative phosphorylation-related proteins were only observed after 20–30 days of PRT. This disconnection between changes in mRNA expression and protein abundance, particularly for mitochondrial proteins, is consistent with findings in humans (Robinson et al., 2017, Tharakan et al., 2021) and warrants further exploration.

In humans, acute increases in mixed-protein FSR after early bouts of resistance exercise do not predict the magnitude of muscle growth that occurs after chronic RT (Mitchell et al., 2014), and in rats activators of translation initiation (i.e. p70s6K phosphorylation) exhibit a greater response in comparison to subsequent gains in muscle growth (Baar and Esser, 1999). We demonstrate a relatively small (∼27%) proportion of newly synthesised protein is retained and contributes to gains in muscle protein abundance in response to RT (Figure 4). This relatively low ‘conversion factor’ indicates a prominent activation of protein degradative processes in Stim muscle, which is consistent with elevations in the expressions of gene associated with phagosomal and lysosomal processes throughout the 30-day period of PRT (Viggars et al., 2023). A high turnover of muscle protein may be necessary to replace damaged proteins and may improve muscle protein homeostasis in exercised muscle (Srisawat et al., 2023). Our data may also suggest gains in muscle mass in response to RT could be further optimised by investigating the effects of manipulating, for example, training variables or nutritional interventions, to increase the ‘conversion factor’ between the synthesis and accretion of muscle protein. In particular, it will be interesting to discover whether age-related losses in muscle mass are associated with a lesser conversion between protein synthesis and accretion.

Investigating the processes of skeletal muscle adaptation to RT requires multifactorial experimental designs, which are currently uncommon in ‘omics studies. We highlight the reliability of the unilateral PRT model and the Proteo-ADPT workflow, evidenced by the high reproducibility between Day 0 Ctrl and Stim muscles and consistent abundance profile of Ctrl muscle across the 30-day experimental period. These technical strengths, combined with stringent data handling enabled us to investigate interaction (Time*Condition) effects, which are not captured by more commonly applied single-factor comparisons. However, our stringent statistical criteria, meant a relatively modest number of proteins were included in the final analysis compared to recent studies investigating only the relative abundance profile of muscle proteins. Nevertheless, our analyses encompass the majority (>70 %) of the total protein content of muscle and were sufficient to distinguish different temporal responses between subsets of ribosomal and mitochondrial proteins. The mitochondrial protein response may be particular to our model of daily PRT, and further work is required to investigate muscle responses to training patterns involving less frequent muscle stimulation. Our programme of daily PRT is known (Viggars et al., 2023) to result in complete silencing of fast-twitch myosin heavy chain, Myh4, mRNA expression and so likely exaggerates mitochondrial adaptations compared to typical human RT interventions that specifically increase the synthesis of fast-twitch IIa myosin heavy chains (Camera et al., 2017).

## Conclusion

Programmed resistance training (PRT) resulted in significant muscle growth, driven by time-dependent changes in the synthesis and abundance of specific proteins. Ribosomal protein accretion and enhanced translational capacity were prominent during the first 10 days of PRT and preceded the later accretion of mitochondrial proteins. These findings demonstrate that muscle hypertrophy in response to PRT involves the sequential accumulation of distinct subsets of the muscle proteome rather than uniform and simultaneous gains across all muscle proteins. Our observations highlight the challenges of studying adaptation without stable isotope labelling and the collection of time-series data or going beyond bulk measurements of mixed-protein synthesis rates to investigate individual protein dynamics and protein accretion in absolute terms. In particular, further exploitation of Proteo-ADPT could help discover new interventions for improving muscle mass and function by optimising protein accretion in addition to protein synthesis.

## Data availability

The mass spectrometry proteomics data have been deposited to the ProteomeXchange Consortium via the PRIDE partner repository with the dataset identifier PXD060879 and 10.6019/PXD060879.

## Author contributions

JC Jarvis and JG Burniston conceived and designed the research; CA Stead, SJ Hesketh, ACQ Thomas, H Sutherland, JC Jarvis, and JG Burniston performed the research and acquired the data. CA Stead, ACQ Thomas, SJ Hesketh, and JG Burniston analysed and interpreted the data. All authors were involved in drafting and revising the manuscript.

